# Impact of mutations in the *HEADING DATE 1* gene on transcription and cell wall composition of rice

**DOI:** 10.1101/2024.10.16.618758

**Authors:** Marco Biancucci, Daniele Chirivì, Alessio Baldini, Eugene Badenhorst, Fabio Dobetti, Bahman Khahani, Elide Formentin, Tenai Eguen, Franziska Turck, John P. Moore, Elahe Tavakol, Stephan Wenkel, Fiorella Lo Schiavo, Ignacio Ezquer, Vittoria Brambilla, David Horner, Matteo Chiara, Giorgio Perrella, Camilla Betti, Fabio Fornara

## Abstract

Plants utilize environmental information to modify their developmental trajectories for optimal survival and reproduction. Over a century ago, day length (photoperiod) was identified as a major factor influencing developmental transitions, particularly the shift from vegetative to reproductive growth. In rice, exposure to day lengths shorter than a critical threshold accelerates flowering, while longer days inhibit this process. This response is mediated by HEADING DATE 1 (Hd1), a zinc finger transcription factor that is central in the photoperiodic flowering network. Hd1 acts as a repressor of flowering under long days but functions as a promoter of flowering under short days. However, the transcriptional organization of this dual function is still not fully understood. In this study, we utilized RNA-Seq to analyze the transcriptome of *hd1* mutants under both long and short day conditions. We identified genes involved in the phenylpropanoid pathway that are deregulated under long days in the mutant. Quantitative profiling of cell wall components and abiotic stress assays suggest that Hd1 is involved in processes considered unrelated to flowering control. This indicates that day length perception and responses are intertwined with physiological processes beyond flowering.

## Introduction

The timing of flowering is an important adaptive trait for all plant species. It allows synchronization of the reproductive phase with optimal seasonal conditions and among individuals, thus maximizing seed set. This feature is particularly relevant for crop species because it sets cycle length, ensures maximal yields, and facilitates field management. The trait is under tight genetic and environmental control, and a very large number of flowering time genes, arranged in regulatory networks, work at the interface between monitoring endogenous and external parameters and promoting or repressing flower development.

Several factors can influence seasonal flowering, including aging and hormones, water and nutrient availability, biotic and abiotic stresses, fluctuating temperatures and light conditions (Song et al., 2015; Vicentini et al., 2023). However, among all parameters, changes in day length (photoperiod) are the most informative because their pattern is invariant from year to year and therefore predictable and reliable for anticipating seasonal changes. Plants have evolved the capacity to measure and respond to day length variations and can be categorized as long (LD) or short day (SD) species, depending on the condition that promotes flowering. Day neutral behaviors are also observed, wherein species do not use the photoperiod as an environmental cue to control flowering.

Rice is a facultative SD plant, in which flowering is accelerated when the photoperiod falls below a critical threshold (Itoh et al., 2010). Its progenitors can be found in tropical and subtropical regions (Wang et al., 2018; Jing et al., 2023). However, breeding efforts have succeeded in expanding cultivation also to higher latitudes, characterized by LD during the cropping season, in both Asia and Europe (Gómez-Ariza et al., 2015; Goretti et al., 2017; Zong et al., 2021; Sun et al., 2022).

A complex regulatory network, principally constituted by photoreceptors and transcription factors, measures day length, and determines flowering time. The *HEADING DATE 1* (*Hd1*) gene was the first component of the rice photoperiodic network to be cloned and is a close homolog of *CONSTANS* (*CO*), a photoperiod sensor of Arabidopsis (Yano et al., 2000). Both genes encode transcription factors characterized by the presence of B-Box zinc finger domains at the N-terminus and of a CONSTANS, CO-like, and TOC1 (CCT) domain at their C-terminus, which are required for protein-protein interactions and DNA binding. *Hd1* and *CO* are not orthologs, and their recruitment in the photoperiodic network is likely the result of convergent evolution (Ballerini and Kramer, 2011; Simon et al., 2015; Vicentini et al., 2023). Functionally, *Hd1* promotes flowering under SD, by inducing transcription of *HEADING DATE 3a* (*Hd3a*) and *RICE FLOWERING LOCUS T 1* (*RFT1*), encoding rice florigens. Its activity reverts under LD, and *Hd1* becomes a repressor of flowering and of *Hd3a* and *RFT1* expression. *CO* shows a similar photoperiod-dependent functional reversion, promoting flowering under LD and repressing it under SD, although its SD repressive activity is not dependent upon reduction of florigen expression (Luccioni et al., 2019).

The Hd1 protein forms higher-order heterotrimeric NUCLEAR FACTOR Y (NF-Y) complexes, interacting with NF-YB and NF-YC subunits (Goretti et al., 2017; Shen et al., 2020). This feature is typical of proteins containing a CCT domain and occurs among both monocot and dicot species (Wenkel et al., 2006; Li et al., 2011; Goretti et al., 2017; Shen et al., 2020). NF-YB and NF-YC form histone-like dimers that have non-sequence specific affinity for DNA. When a CCT domain protein is incorporated, the trimer binds specifically to sequences containing a CO-Responsive Element (*CORE*). Initial studies on Arabidopsis defined the *CORE* as *TGTG(N2-3)ATG* (Wenkel et al., 2006; Adrian et al., 2010; Tiwari et al., 2010; Gnesutta et al., 2017).

Subsequent work narrowed down the *CORE* to *TGTGGT* (for potato StCOL1) and *TGTGG* (for Arabidopsis CO and rice Hd1) (Abelenda et al., 2016; Goretti et al., 2017; Gnesutta et al., 2017). The crystal structures of CO and Hd1, in complex with NF-YB/C subunits and DNA, have further refined the *CORE*, indicating that essential contacts are made with a *TGTG* motif only (Shen et al., 2020; Chaves-Sanjuan et al., 2021; Lv et al., 2021).

Rice NF-Y can accommodate distinct DNA binding subunits containing a CCT domain. These include GRAIN YIELD PLANT HEIGHT AND HEADING DATE 7 (Ghd7), PSEUDO RESPONSE REGULATOR 37 (PRR37) and PRR73 (Shen et al., 2020; Liang et al., 2021). Changing the DNA binding subunit might modify preference of the trimer for motifs recognition. However, comparison of available binding motifs, identified by chromatin immuno-precipitation, SELEX, or other techniques, suggests that all CCT domain proteins might bind a *TGTG* core sequence (Gnesutta et al., 2018).

The rice NF-YB and NF-YC subunits belong to expanded families, comprising 11 and 7 genes, respectively, indicating a certain degree of redundancy and/or cooperativity (Petroni et al., 2012). The *OsNF-YB11* gene encodes for Ghd8/DTH8/Hd5 (hereafter Ghd8), a major LD repressor in the photoperiod pathway (Wei et al., 2010). The OsNF-YB7, 8, 9 and 10 proteins have similar activities, although not as central as Ghd8, and could replace Ghd8 in the heterotrimer (Hwang et al., 2016; Li et al., 2016). Similarly, biochemical and genetic evidence point to redundant roles for OsNF-YC1, 2, 4 and 7 (Kim et al., 2016; Goretti et al., 2017; Shen et al., 2020). A direct interaction between Hd1 and Ghd7 has also been reported, suggesting that NF-Y complexes might include more than one CCT protein, or that multiple NF-Y complexes interact through their CCT components. Hd1 and Ghd7 repress expression of *EARLY HEADING DATE 1* (*Ehd1*), a central promotor in the flowering network, under LD (Nemoto et al., 2016).

Single mutations in *Ghd7* or *8*, *PRR37*, *PRR73* and several *NF-YC* genes accelerate flowering under LD, consistent with their involvement in LD repressor complexes (Xue et al., 2008; Wei et al., 2010; Koo et al., 2013; Gao et al., 2014; Kim et al., 2016; Liang et al., 2021). In plants harboring *ghd7* or *ghd8* mutations and grown under LD, Hd1 is converted from a repressor to an activator of flowering (Du et al., 2017; Zong et al., 2021; Sun et al., 2022). These genetic data support the hypothesis that the switch in Hd1 function depends upon incorporation of Ghd7 and/or Ghd8 into LD repressor complexes. Under SD inductive conditions, transcription of *Ghd8* is low and Ghd7 protein accumulation is prevented by post-transcriptional mechanisms, thus releasing the promoting activity of Hd1 (Zheng et al., 2019). Whether Hd1 forms different complexes under SD remains to be determined.

The targets of Hd1 include *Hd3a* and *RFT1*, encoding for florigenic proteins expressed in phloem companion cells and loaded into sieve elements. Once in the phloematic stream, they can reach the shoot apical meristem (SAM), acting as long-distance, non-cell autonomous signals and promoting the transition of the apex from vegetative to reproductive. While several lines of evidence support phloematic expression of Hd3a and RFT1, the question of whether Hd1 expression is limited to the phloem is still unanswered (Tamaki et al., 2007; Komiya et al., 2009; Pasriga et al., 2018) It is equally unknown whether Hd1 has additional targets, either direct or indirect.

In this study, we used genome-wide and biochemical approaches to explore the regulatory landscape of the Hd1 protein.

## Results

### Mutations in *Hd1* modify the leaf transcriptome more extensively under LD

In Arabidopsis, *CONSTANS* is transcribed in companion cells of the phloem. Its misexpression under the companion cell-specific *SUCROSE TRANSPORTER 2* (*SUC2*) promoter is sufficient to accelerate flowering, whereas misexpression in the SAM does not result in appreciable changes in flowering time (An et al., 2004). In rice, transcription of *Hd1* has not been studied at the tissue-level, although the gene is also assumed to be expressed in vascular tissues, because *Hd3a* and *RFT1* are activated there (Komiya et al., 2008; Pasriga et al., 2018). We stably transformed a *pHd1:GUS* vector previously used in transient assays (Goretti et al., 2017) into Nipponbare, and analyzed GUS expression patterns of independent T2 transgenic plants under LD conditions (Figure 1A-D). During early developmental stages, when plants were 3 weeks old, GUS expression was detected in the vascular tissue of the leaf (Figure 1A-B). At advanced stages of development, when plants were 6 weeks old, GUS expression was detected in the phloem as well as in all mesophyll cells, but not in the epidermis (Figure 1C-D). This pattern is consistent with *Hd1* controlling expression of *Hd3a* and *RFT1* in the vasculature, but also suggests that *Hd1* has a broader expression, and might target additional genes controlling physiological processes other than flowering time.

**Figure 1.**
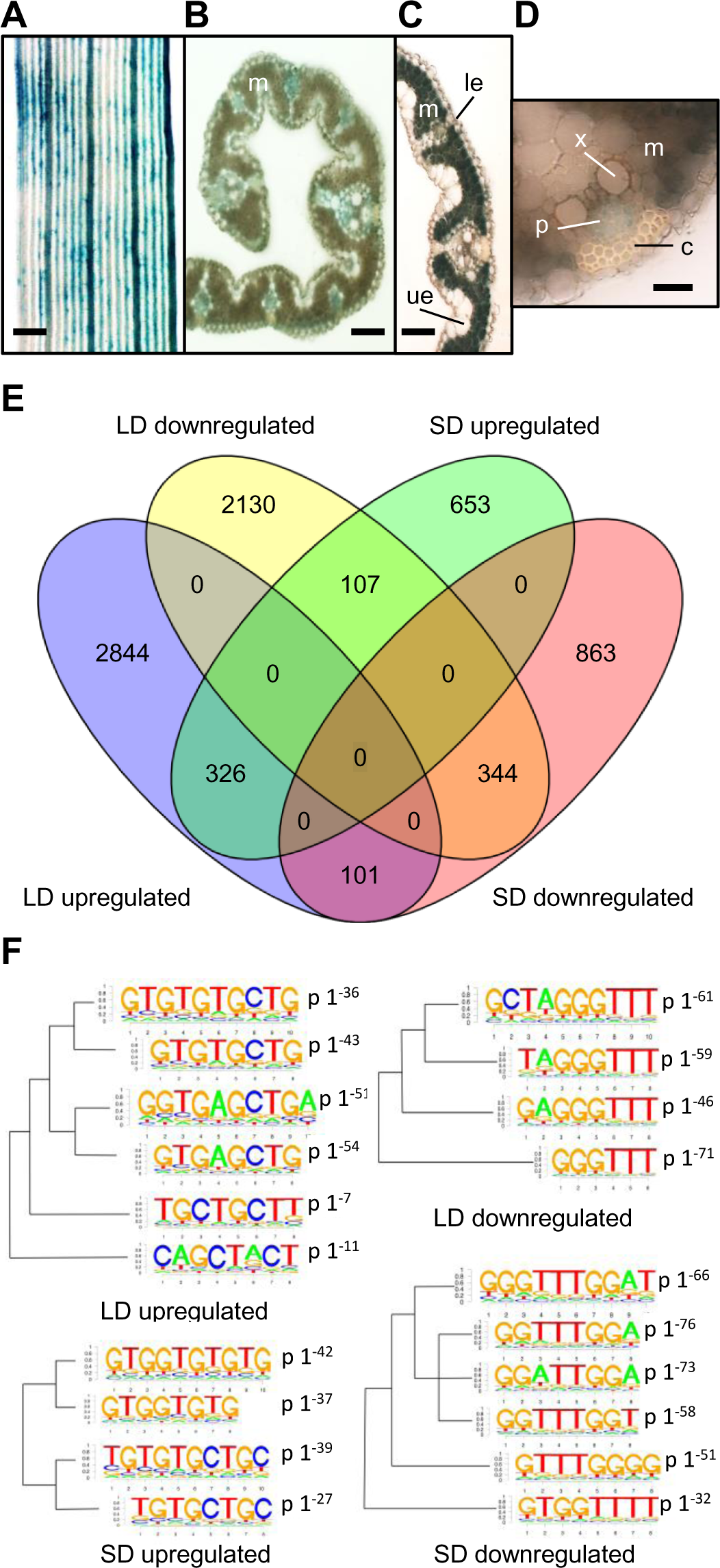
Transcriptional changes caused by *Hd1* in the leaf under LD and SD. **A-D**, GUS assays on rice leaves transformed with a *pHd1:GUS* vector. **A-B**, 3-week-old rice leaves showing GUS expression in the vasculature. **C**, 6-week-old leaf showing GUS expression in the mesophyll. **D**, magnification of a 6-week-old vascular bundle showing details of conductive tissues. Scale bars: A=100µm, B and C=50µm, D=20µm; m, mesophyll; le, lower epidermis; ue, upper epidermis; p, phloem; c, collenchyma; x indicates a vessel element cell of the xylem. **E**, Venn diagram summarizing genes differentially expressed in *hd1-1* compared to wild type, under LD and SD at FDR<0.05. **F**, logo plots of enriched DNA motifs in the promoters of DE genes, filtered for FDR<0.05 and log_2_FC≥|1.5|.

Following this hypothesis, we performed a global analysis of gene expression by RNA-sequencing, comparing the leaf transcriptomes of *hd1-1* mutants vs. Nipponbare wild type, under both LD and SD. Triplicate samples were collected at ZT0, 70 and 56 days after sowing in LD and SD, respectively. Flowering time and the expression of known *Hd1* target genes, including *Hd3a* and *RFT1*, were quantified to assess proper growth conditions and transcription patterns. The results were consistent with published data (Supplementary Figure 1A-C).

When applying an FDR≤0.05, we identified 5852 differentially expressed genes (DEGs) between *hd1-1* mutants and wild-type Nipponbare under LD, with a slight overrepresentation (56%) of upregulated genes (Figure 1E; Supplementary Table 1 and 2 report the complete lists of genes from LD and SD experiments, respectively). Under SD, 2394 genes were differentially expressed, less than half as many as under LD, with a slight overrepresentation (55%) of downregulated genes.

Differences became even more evident when filtering also for fold change (FC). An arbitrary log_2_FC≥|1.5| reduced LD DEGs to 2188, and SD DEGs to 81 only. These data indicate that *Hd1* has a greater effect on the transcriptome under LD than under SD.

The abundance of Hd1 protein cycles during the day and is highest during the light phase. The accumulation profile is the result of translation from cycling RNA, as well as protein degradation - mediated by the autophagy pathway - in the dark (Yang et al., 2015; Hu et al., 2022). With the list of LD DEGs filtered by log_2_FC≥|1.5|, we used Phaser to determine if specific peak expression phases were enriched among the clock-controlled genes whose expression also depends upon Hd1 (Mockler et al., 2007). The data indicated that, of the cycling genes, those having peak expression at ZT0-1 and ZT4-9 were enriched in the dataset, with respect to random sampling (Supplementary Figure 1D). These observations are consistent with the hypothesis that mutations in *Hd1* have a stronger impact on genes with peaks of expression occurring during the light.

Next, we compared genes differentially expressed in *hd1* under either LD or SD, with genes whose expression depends upon the shift from long to short day lengths, using datasets in which conditions were very similar (albeit not identical) to those used in this study (Galbiati et al., 2016). The scope of this meta-analysis was to quantify the overlap between the *hd1*- and photoperiod-dependent transcriptomes, and possibly identify overrepresented categories at their intersection. Only 8 genes were in common to the *hd1* SD and photoperiod datasets (but not *hd1* LD), and among them *OsMADS1*, *OsMADS14* and *Hd3a* were identified as being transcribed in response to *Hd1* and under SD (Supplementary Figure 2). Thus, while short, this list contains genes with physiological roles during the reproductive phase. The function of *OsMADS1* and *OsMADS14* in leaves is still unclear, although the latter is known to be expressed only under SD, with a peak of expression occurring during the night (Brambilla et al., 2017). The overlap between genes regulated by *Hd1* under LD and those controlled by photoperiod consisted of 403 genes. This overlap is highly significant by factor of over-representation (4) and p value (p<3.1 e-136).

However, we did not find enriched functional categories within this group. Thus, the LD transcriptome of *hd1* shared similarities with that of SD-treated wild type plants, but no specific pathway or functional category was evident.

Finally, we retrieved the promoters of DEGs spanning-1Kb to +100bp from the transcriptional start site (TSS) and scanned them with algorithms for *de novo* discovery of binding sites based on motifs enrichment (see Methods). Among promoters of DEGs, we identified several motifs statistically supported, both among up and downregulated genes, in both photoperiods (Figure 1F). Interestingly, promoters of upregulated genes frequently harbored a *TGTG* sequence, which is also present in *FT* and *Hd3a CORE* regions, and essential for binding of CO/NF-Y and Hd1/NF-Y, respectively. Promoters of downregulated genes were enriched with sequences containing *GGTTT*. The difference between enriched motifs did not depend on day length, but on the direction of differential expression, indicating an *Hd1*-dependent effect. This analysis does not demonstrate direct binding of Hd1 to enriched motifs. One possibility is that Hd1 changes preference for DNA, depending on whether it acts as promotor or repressor of transcription. However, it should be noted that, given the reduced depth of the SD transcriptome data, *GGTTT* motifs require further validation.

### Genes belonging to the phenylpropanoid pathway are enriched among DEGS in the *hd1* LD transcriptome

Next, we determined enrichment of specific categories by performing Gene Ontology (GO) analyses (Supplementary Figure 3 shows ontology groups statistically enriched according to the Gene Ontology Resource database, https://geneontology.org).

Under LD, we observed several GO terms related to the phenylpropanoid biosynthetic pathway. Many genes encoding enzymes of the pathway were upregulated, consistent with *Hd1* acting as transcriptional repressor (Supplementary Figure 4). To investigate whether the dual transcriptional effect of *Hd1* applied to genes other than *Hd3a* and *RFT1*, we sought phenylpropanoid pathway genes downregulated in *hd1-1* under SD. Among those that were upregulated under LD and downregulated under SD in the RNA-Seq experiment, we selected *PHENYLALANINE AMMONIA LYASE 4* (*OsPAL4*, LOC_Os02g41680). *CINNAMYL-ALCOHOL DEHYDROGENASE* (*OsCAD8B*, LOC_ Os09g23540) was chosen as example of a gene downregulated under LD, to assess if *Hd1* could also act as LD activator of gene expression. We quantified their transcription during 24h time course experiments, expanding on the initial single-time settings of RNA-Seq experiments (Figure 2A-D). We observed reduction of *OsPAL4* expression in SD, while steady-state mRNA levels increased under LD in *hd1-1* at all time points (Figure 2A, C). *OsCAD8B* expression was lower in the mutant under LD, but identical to the wild type under SD, throughout the time course (Figure 2B, D).

**Figure 2.**
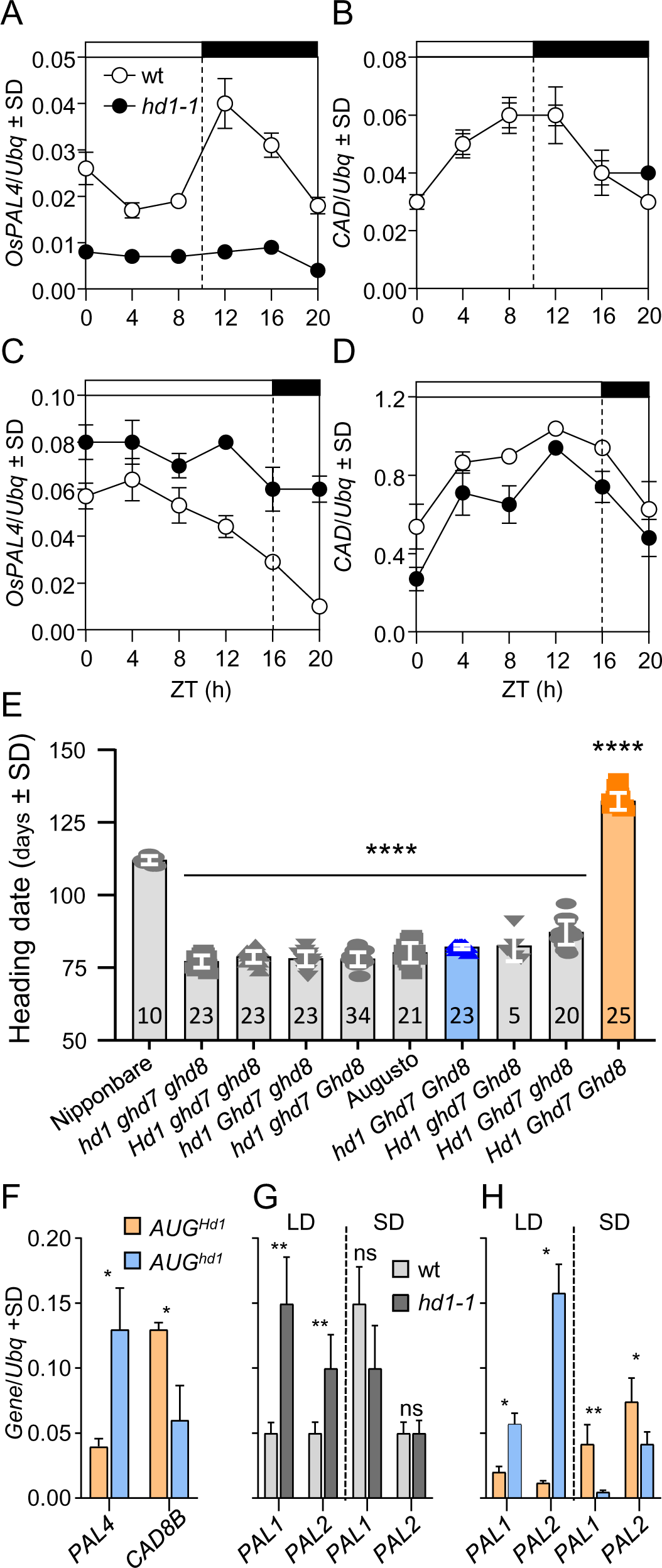
Transcription of genes in the phenylpropanoid pathway. Transcription of *OsPAL4* (**A**, **C**) and *OsCAD8B* (**B**, **D**) quantified under SD (**A**, **B**) and LD (**C**, **D**) in Nipponbare and *hd1-1*. White and black bars on top of the graphs indicate day and night periods, respectively. ZT, *Zeitgeber*. **E**, flowering time of BC3F3 lines scored under natural LD in Milan. The number of plants scored is indicated in each histogram. Genotypes are indicated on the x axis. ****, p<0.0001 based on ordinary one-way ANOVA. **F**, quantification of *OsPAL4* and *OsCAD8B* transcription in field-grown plants harvested 4h after dawn at the summer solstice. **G-H**, quantification of *OsPAL1* and *OsPAL2* transcription in *hd1* mutant alleles under controlled LD and SD. Each quantification represents the average ± standard deviation (SD) of three technical replicates. *UBIQUITIN* (*Ubq*) was used to normalize gene expression. Asterisks indicate statistical significance based on Student’s *t* test. *, p<0.05; **, p<0.005.

We next determined patterns of gene expression in a second *hd1* mutant allele from a different rice variety. To this end, we exploited BC3F3 lines obtained from a cross between Nipponbare and Augusto, with Augusto used as recurrent parent (see Materials and Methods). Augusto harbors loss-of-function alleles of *Hd1*, *Ghd7* and *Ghd8*. The Augusto *hd1* allele has a frameshift mutation that disrupts the CCT domain (Gómez-Ariza et al., 2015). We derived all combinations of *hd1*, *ghd7* and *ghd8* mutations in the Augusto background and used two introgressions selected to bear *hd1^AUG^ Ghd7^NB^ Ghd8^NB^* (hereafter *AUG^hd1^*) and *Hd1^NB^ Ghd7^NB^ Ghd8^NB^* (hereafter *AUG^Hd1^*). As expected, flowering was accelerated in *AUG^hd1^* compared to *AUG^Hd1^*(Figure 2E). Expression of *OsPAL4* and *OsCAD8B* was quantified in leaves of plants grown in a field and harvested at the summer solstice when day length was at its maximum (15h 40m). *OsPAL4* and *OsCAD8B* transcription showed opposite regulation in *AUG^hd1^*compared to *AUG^Hd1^*, consistent with data obtained from controlled growth conditions (Figure 2F).

*OsPAL4* is found in a genomic cluster containing four *PAL* genes, all of which were upregulated under LD in *hd1-1*, based on RNA-Seq data (Supplementary Table 1). We quantified transcription of *OsPAL1* (LOC_Os02g41630) and *OsPAL2* (LOC_Os02g41650) under LD and SD in *hd1-1* and *AUG^hd1^* mutant backgrounds and observed patterns like *OsPAL4* (Figure 2F, G). Quantification of *OsPAL3* (LOC_Os02g41670) mRNA expression failed due to amplification of multiple transcripts in qPCR experiments.

These data indicate that *Hd1* has opposite effects on transcription of *OsPAL4*, similar to the regulation of florigens and that it can also operate on genes not belonging to the photoperiodic flowering pathway.

### Hd1 binds the promoter of *OsPAL4*

We then used chromatin immunoprecipitation (ChIP) to assess binding of Hd1 to DNA. To this end, we exploited a line overexpressing FLAG-tagged Hd1 under the control of the maize *ACTIN* promoter (*pACT:3xFLAG:Hd1*). Plants harbouring this vector produce a FLAG-Hd1 protein of the expected size and flower late under 16.5h photoperiods (Eguen et al., 2020). We measured FLAG-Hd1 protein abundance at several time points, under the same growth conditions used for the LD RNA-Seq experiment and observed similar accumulation at every time of day tested (Figure 3B). The protein accumulation pattern followed the transcriptional pattern (Figure 3A), and no evidence of post-translational control of protein abundance was evident, although it is possible that high protein expression might have masked regulatory layers relevant in a wild type context. Nonetheless, this experiment indicated that the Hd1 protein is stable *in vivo*, distinct from the situation observed for CO. We used leaves harvested at ZT1 for chromatin preparations. Following IP with anti-FLAG antibodies, we quantified DNA at the *Hd3a* and *OsPAL4* promoters. The *OsCORE2* motif in the *Hd3a* promoter was used as a positive control, because it has been previously assayed for Hd1 binding *in vitro* (Goretti et al., 2017). We observed Hd1 binding at the *OsPAL4* locus, in a region spanning several *TGTGG* motifs (Figure 3C, D). Therefore, Hd1 directly binds the *OsPAL4* promoter and regulates its transcription, similarly to *Hd3a*.

**Figure 3.**
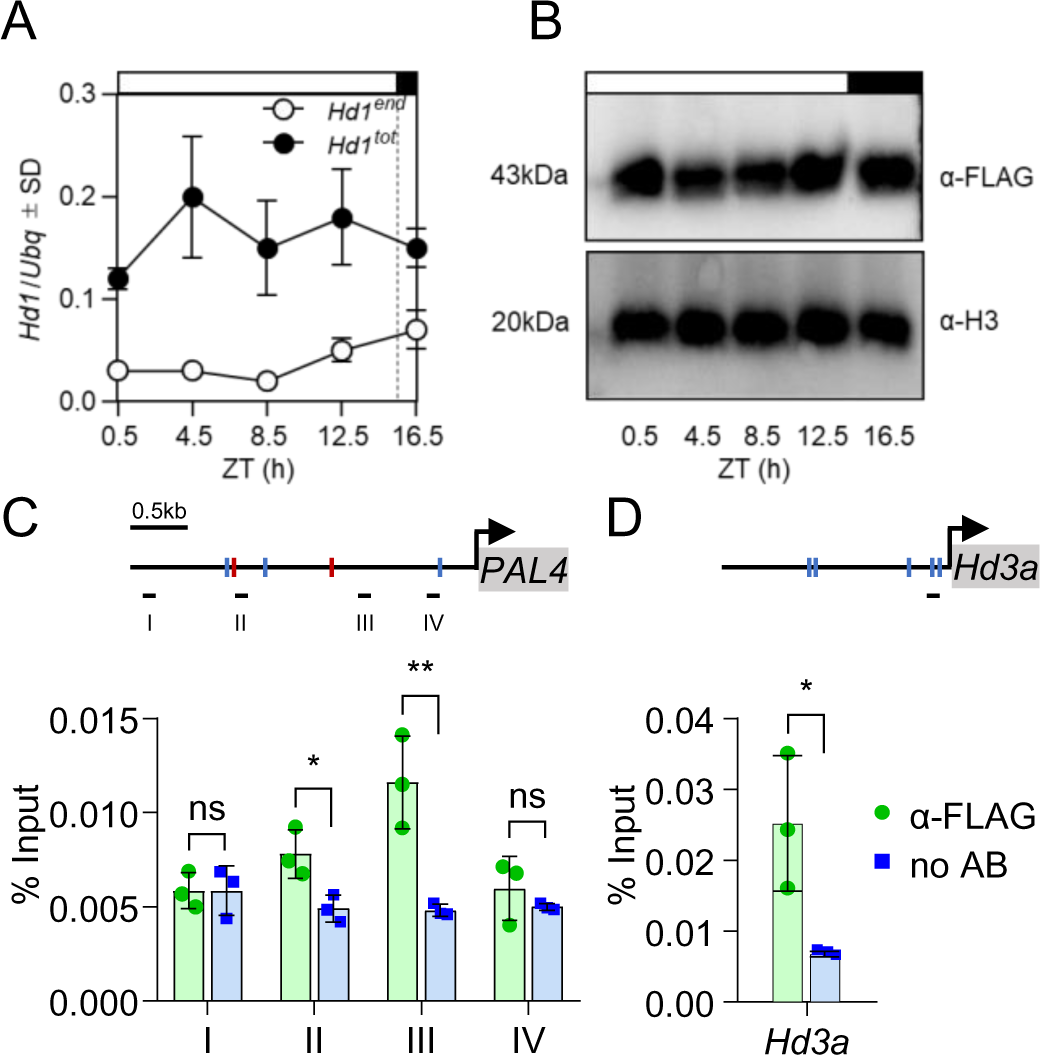
Hd1 binds the *OsPAL4* promoter. Diurnal accumulation profile of endogenous (*Hd1^end^*) and endogenous + transgenic *Hd1* (*Hd1^tot^*) mRNA from a time course in leaves under LD (**A**), compared to accumulation of 3xFLAG-Hd1 from the same samples (**B**). Western blots were repeated twice with biologically independent samples, giving the same results. Anti-histone H3 was used as loading control. Transcriptional quantifications represent the average ± standard deviation (SD) of three technical replicates. *UBIQUITIN* (*Ubq*) was used to normalize gene expression. ZT, *Zeitgeber*. ChIP-qPCR quantifications of Hd1 binding to the promoter regions of *OsPAL4* (**C**) and *Hd3a* (**D**). Schemes on top of the graphs indicate the promoter regions. Red and blue marks indicate *TGTGG* motifs on the plus and minus strands, respectively. Black lines below the promoters indicate the position of the amplicons used to quantify fragments enrichment. Each bar represents the average ± standard deviation (SD) of three technical replicates. Values are shown relative to the input. ChIP-qPCRs were repeated four times independently, giving the same results. Asterisks indicate statistical significance based on Student’s *t* test. *, p<0.05; **, p<0.005.

### The leaf proteome is modified by changes in day length

We asked how day length and/or Hd1 might alter the leaf proteome. Therefore, we carried out total leaf proteome analysis in the Nipponbare wild type and *hd1-1* plants under both SD and LD conditions. Triplicate samples were collected at ZT0 for each condition. Leaves were harvested 30 days after sowing (LD) and after 15 additional days of growth under SD. Mass spectrometry was performed for untargeted proteomics. A total of 6186 proteins were identified. Comparisons between photoperiods showed that 283 were significantly enriched under LD and 311 were significantly enriched under SD conditions in the wild type. In the *hd1-1* mutant, the equivalent numbers were 474 under LD and 462 under SD conditions (Supplementary Figure 5 and Supplementary Table 3). Comparisons between genotypes under the same photoperiodic conditions showed negligible differences under SDs (1 protein more abundant in the wild type and 1 in *hd1-1*). Under LD conditions, 19 proteins were more abundant in the wild type and 12 in the *hd1-1* mutant. We attribute these marginal differences between genotypes to the depth of total proteome analyses, which likely capture only major differences. Nevertheless, these data indicate that changes in daylength have a prominent effect on the leaf proteome, and that changes during the photoperiodic transition are accentuated by the *hd1* mutation. Analysis of ontological categories indicated that changes in day length affected several metabolic processes. On the contrary, comparison between wt and *hd1-1* under LD conditions identified only GO terms related to cell wall metabolism (Supplementary Figure 6).

### Mutations in *Hd1* modify cell wall composition

Both RNA and protein profiling suggest that Hd1 could affect cell wall composition and biogenesis. Therefore, we performed a more detailed biochemical characterization. To this end, we extracted cell wall polymers from the alcohol insoluble residue of *hd1-1* and wild type, using either 50 mM cyclohexane-1,2-diaminetetraacetic acid (CDTA) or 4M sodium hydroxide (NaOH). CDTA at relatively low concentration solubilizes polymers with weak association to the cell wall, whereas NaOH solubilizes polymers strongly attached to it (Ezquer et al., 2016). We will refer to the CDTA and NaOH extractions as soft and harsh, respectively. Quantifications were performed by enzyme-linked immunosorbent assay (ELISA), using the set of antibodies listed in Supplementary Table 4.

Pectins are typically soluble in water or CDTA. However, the harsh treatment released extra material that was not extracted with the soft treatment. In the soft extract, *hd1-1* exhibited significantly lower signals for the backbone of rhamnogalacturonan I (RG-I, backbone of alternating galacturonic acid and rhamnose and typical side chains consisting of arabinose and galactose; p<0.01) and unesterified homogalacturonan (HG, linear chain of galacturonic acid to which methyl or acetyl groups can be attached; p<0.05) when compared to the wild type (Figure 4A, C). In contrast, in the harsh extract, *hd1-1* showed increased abundance of both RG-I side chains, β-1-4-galactan (p<0.01) and α-1-5-galactan (p<0.01) (Figure 4B, D). These data suggest that pectins belonging to the RG-I group are more ramified in the *hd1-1* mutant.

**Figure 4.**
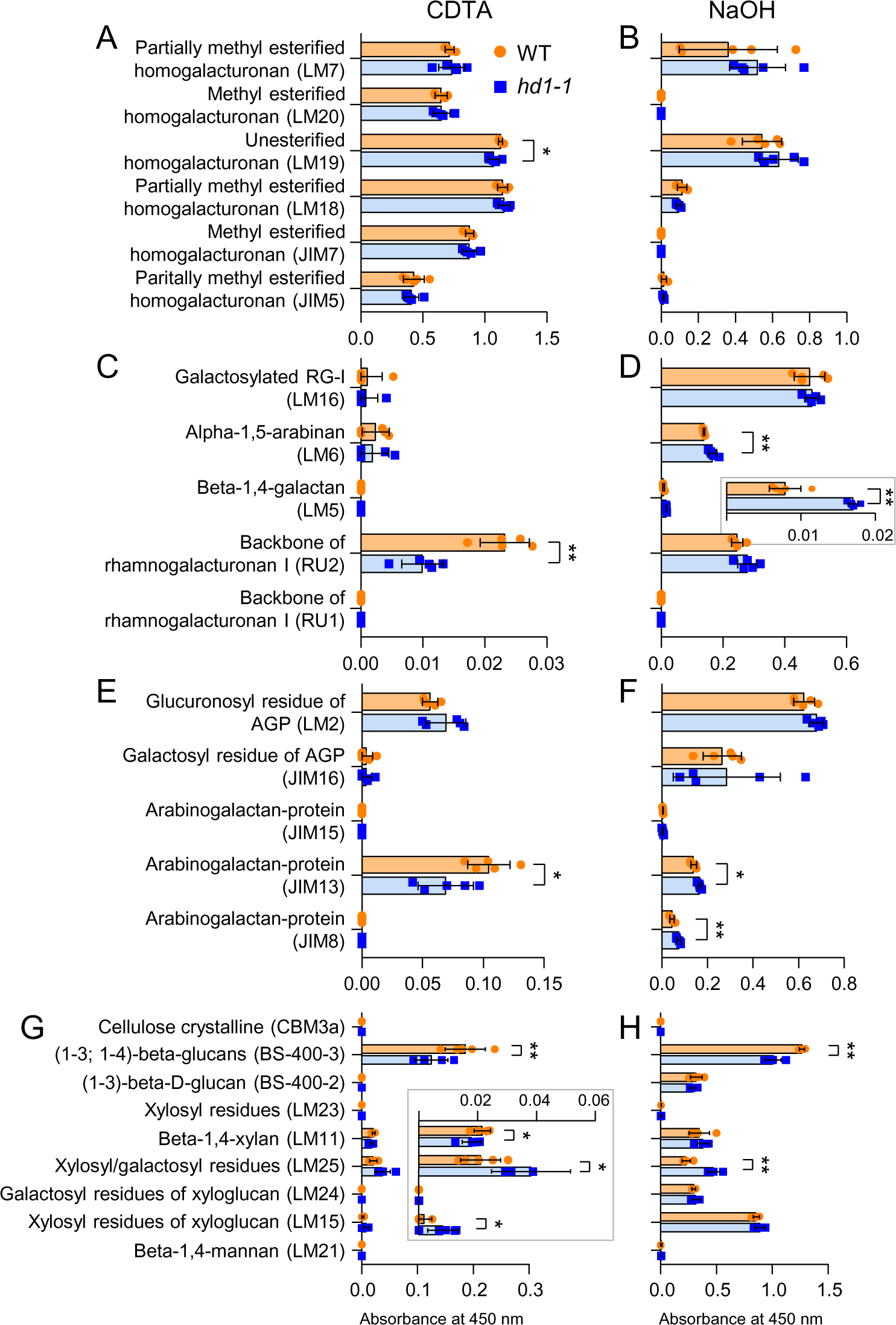
Cell wall composition of the *hd1* mutant. **A**, **C**, **E**, **G**, quantifications of loosely adhered components (CDTA extractions). **B**, **D**, **F**, **H**, quantifications of strongly adhered components (NaOH extractions). **A-D**, quantifications of pectins. Histograms are divided into two groups (**A**, **C** and **B**, **D**) to facilitate reading, because of the different scales of values. Inset in D magnifies the corresponding beta-1,4-galactan values. **E-F**, quantifications of arabinogalactan proteins. **G-H**, quantifications of crystalline cellulose and hemicellulose. Inset in **G** magnifies the corresponding beta-1,4-xylan, xylosyl/galactosyl residues and xylosyl residues of xyloglucan values. Bars indicate the average ± standard deviation of five biological replicates, except for beta-1,4-mannan values where 3 and 4 replicates were used for *hd1-1* and wt, respectively. Each dot represents an independent sample. Asterisks indicate statistical significance based on two-tailed Student’s *t* test. *, p<0.05; **, p<0.01.

Arabinogalactan-proteins (AGP) are highly glycosylated proteins integral to plant cell walls and involved in many activities related to cell growth and development. We used five antibodies to profile AGPs of cell wall preparations. In the soft extract, *hd1-1* showed lower abundance of AGPs detected by JIM13 (recognizing carbohydrate residues of AGPs located on the outer surface of the plasma membrane; p<0.05, Figure 4E). In the harsh extraction, JIM8 and JIM13 produced stronger signals in the *hd1-1* mutant compared to the wild type (Figure 4F). JIM8 has similar properties as JIM13, but it immunoreacts with less AGPs. This may indicate that AGPs that strongly adhere to the extracellular matrix are more prevalent in the mutant. However, given the large number of secreted AGPs and their polymorphisms, this conclusion requires further support.

The fibrillar component of the cell wall was assayed using antibodies marking cellulose as well as some epitopes for hemicellulose. The *hd1-1* mutant had lower levels of (1-3; 1-4)-β-glucans in both the CDTA (p <0.01; Figure 4F) and NaOH (p<0.01; Figure 4G) extractions. Lower signal intensities were also detected for β-(1-4)-xylan, although only in soft extractions and at marginal statistical significance (p<0.05; Figure 4F). Xyloglucans are composed of variable building blocks, formed by linear glucans to which xylosyl and galactose units can be added. Building blocks made of four glucosyl units, α1,6-linked to three xylosyl units, are indicated as XXXG. in turn, xylosyl units can be β1,2-linked to one or two galactose units, and are indicated as XXLG and XLLG, respectively. The *hd1-1* mutant showed higher levels of xylosyl/galactosyl residues (XXLG and XLLG motifs) both in the soft (p<0.05; Figure 4F) and harsh (p<0.01; Figure 4G) extractions. It also showed higher levels of XXXG motifs, detected by LM15, but only in CDTA extractions at p<0.05 (Figure 4F). These data indicate that *hd1* mutations alter the fibrillar component of leaves cell walls, reducing (1-3; 1-4)-β-glucans and β-(1-4)-xylan, while increasing xyloglucans of the XXLG and XLLG types.

### Mutations in *Hd1* change salt stress tolerance in a photoperiod-dependent manner

The results presented so far indicate that *Hd1* could have broader roles than the control of flowering time and affect other physiological processes. We hypothesised that abiotic stress tolerance might be altered in the mutant, also considering that GO categories suggested involvement in response to external stimuli and to abscisic acid - a stress hormone - detoxification of reactive oxygen species and general defence responses (Supplementary Figure 3). We choose salinity stress to challenge this hypothesis. We grew wild type and *hd1* mutants in artificial media containing 300 mM sodium chloride, and measured shoot growth as proxy of salt sensitivity, calculating an index based on comparison between treated and non-treated plants (see Materials and methods section). We observed that salt sensitivity did not change in the wild type grown under different day lengths. However, in *hd1-1* and *hd1-2* mutant alleles, salt sensitivity diverged depending on the photoperiod, increasing under LD and decreasing under SD (Figure 5A). Thus, despite no statistically significant difference was observed between wild type and mutant plants grown in the same photoperiod, salt stress was perceived differently by *hd1* mutant plants grown in LD and SD. We assayed *pACT:3xFLAG:Hd1* and observed no difference between photoperiods, but significant reduction of sensitivity compared to *hd1* mutants under LD (Figure 5A). In Augusto, salt sensitivity was less variable compared to Nipponbare, in which ample variability was evident, particularly under SD (Figure 5B). Yet, also in this variety, the *AUG^hd1^*genotype showed differential sensitivity to salt stress, depending on day length. Finally, we assayed an introgression harboring *Hd1^NB^ ghd7^AUG^ ghd8^AUG^* (hereafter *AUG^ghd7,8^*), whose flowering time was very similar to that of *AUG^hd1^*, having loss-of-function alleles of *Ghd7* and *Ghd8* LD repressors (Figure 2E). Salt sensitivity of *AUG^ghd7,8^* diverged similarly to *hd1* mutants across photoperiods, despite marginal statistical significance (Figure 5B). Taken together, these data indicate that rice plants respond differently to salt stress, depending on the photoperiod, but only in genetic backgrounds in which LD floral repression is relaxed.

**Figure 5.**
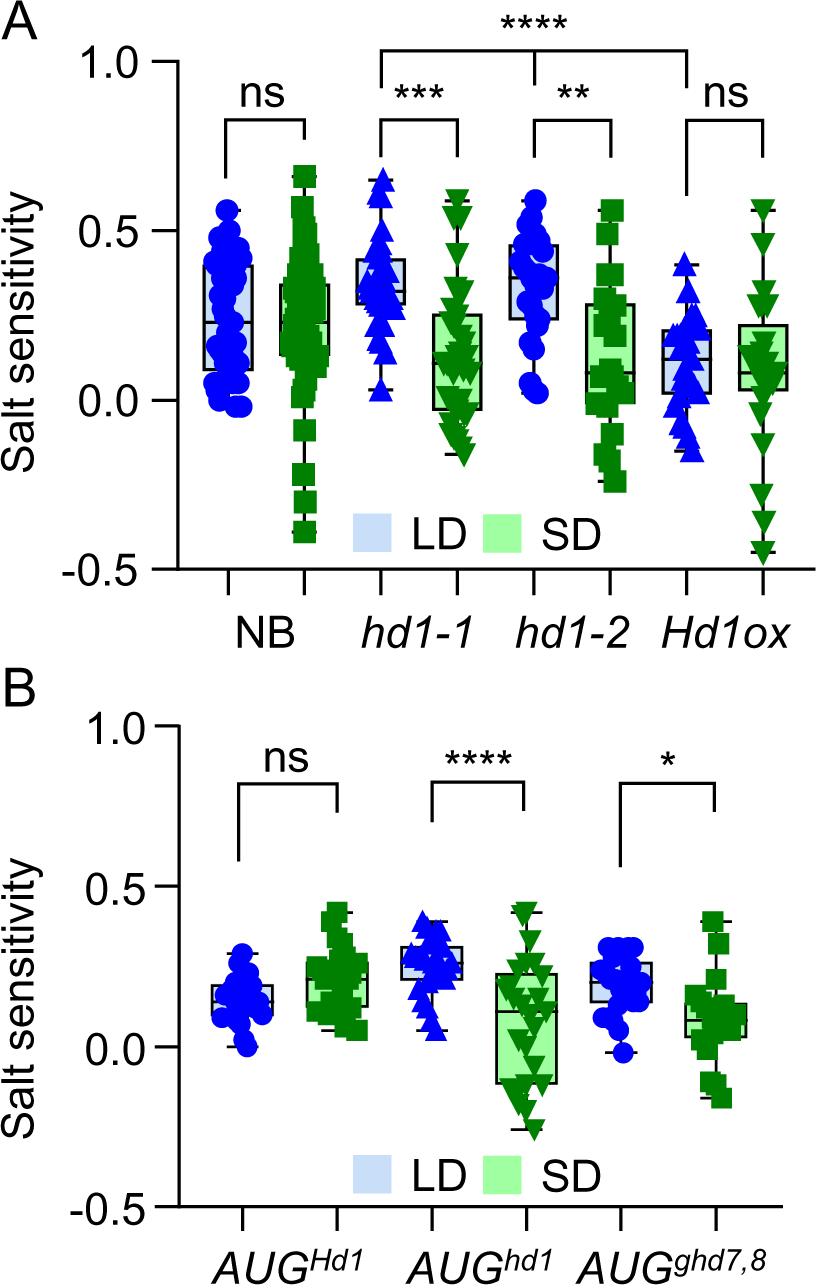
Salt stress assays in *hd1* mutants. **A**, box plots showing salt sensitivity of Nipponbare, *hd1-1*, *hd1-2* and *pACT:3xFLAG:Hd1* (*Hd1ox*). **B**, box plots showing salt sensitivity of Augusto introgression lines. Each box indicates the 25th–75th percentiles, the central line indicates the median and the whiskers indicate the full data range. Each dot indicates a pair of plants (control and treated) used to calculate the index. Pairs of measurements were randomly sampled and used only once. **, p<0.05; ***, p<0.005; ****, p<0.0001 based on ordinary one-way ANOVA. The experiment was repeated three times independently, with similar results.

## Discussion

Plants experience continuous changes in day length, even at latitudes close to the equator, and have adapted to anticipate and respond to them. Flowering time is very susceptible to such changes, and observation of the flowering behaviour of certain species has been instrumental to the recognition of photoperiod measurement systems (Garner and Allard, 1920). However, recent studies have demonstrated that changes in photoperiod can influence several processes unrelated to flowering, including bud dormancy in trees, tuber or bulb formation, and growth, to mention a few important examples (Lee et al., 2013; Abelenda et al., 2016; Tylewicz et al., 2018; Wang et al., 2024). Thus, the photoperiodic pathway, originally and commonly studied in the context of flowering, can be integrated in broader response systems. In rice, Hd1 is central in the photoperiodic flowering pathway and when mutated, alters the capacity of the plant to correctly perceive seasonal changes and flower at the correct time of the year. The data presented in this study, extend the roles of Hd1 and suggest that it has a broader impact on plant physiology.

### Cell wall remodelling in the *hd1* mutant

We have found links between *Hd1* and genes controlling secondary metabolism and cell wall biogenesis, under LD. This observation could be interpreted by postulating either a direct effect of Hd1 on these pathways, or an indirect effect caused by reduced day length sensitivity of the *hd1* mutant. Binding of Hd1 to the promoter of *OsPAL4* supports the former hypothesis, but both could be valid, and more thorough analyses are required to distinguish between them. Nevertheless, quantification of cell wall polymers detected differences between wt and mutant, allowing to draw conclusions on the role of Hd1 in cell wall remodelling.

The cell wall is a highly organized structure enclosing every plant cell, and formed by polysaccharides, proteins and phenolic compounds (Cosgrove, 2024).

Polysaccharides include cellulose, hemicellulose and pectin. Cellulose is a homopolymer of β-(1,4)-D-glucose, and the main component of primary walls. Hemicellulose includes a heterogenous group of polysaccharides formed by a backbone of 1,4-beta-linked sugars, to which side chains of 1-3 sugar residues are covalently linked. The most common hemicellulose of flowering plants is xyloglucan (XyG). However, the cell wall of grasses contains small amounts of XyG, and the most abundant hemicellulose is arabinoxylan. Pectin is found mainly on the outer side of the wall, in the middle lamella, working as a glue between cells. The building block of pectin is α-(1–4)-D-galacturonic acid, forming homopolymers (homogalacturonan) or heteropolymers of rhamnose and galacturonan (RG-I). Homogalacturonan can also present rhamnogalacturonan side chains (RG-II), as well as xylose or other monosaccharide substitutions.

Cellulose is polymerized by CELLULOSE SYNTHASEs (CESA), that reside on the plasma membrane and are organized in multimeric cellulose synthase complexes (CSCs). Cellulose biosynthesis follows diurnal oscillations depending on light and carbon availability, but not on the circadian clock (Ivakov et al., 2017). Seasonal photoperiodic patterns in cellulose biosynthesis have also been observed. In Arabidopsis, the blue-light photoreceptor FLAVIN-BINDING KELCH REPEAT, F-BOX 1 (FKF1) stabilizes CO to promote flowering under LD, while inhibiting cellulose biosynthesis in the leaves (Yuan et al., 2019), providing direct evidence of the connection between the photoperiod pathway and cellulose production.

Both extraction profiles indicated that *hd1* contains less (1-3; 1-4)-beta-glucans and more xylosyl residues of XyG (especially XXLG and XLLG types) in mature leaves. Since the (1-4)-beta-glucan backbone is common to both cellulose and XyG, these data suggest that in *hd1* cellulose is less abundant, XyG are shorter and more ramified, or both. In wt rice, *OsCESA3* and *6* are ubiquitously expressed and are necessary to synthetize microfibrils in primary walls. Expression of *OsCESA4*, *7* and *9* has also been detected in most rice tissues at relatively high levels, with the notable exception of mature leaves, in which transcript abundance is very low or undetectable (Tanaka et al., 2003; Wang et al., 2010). Among the DEGs, *OsCESA1*, *4*, *6*, *7*, *9* and several *OsCESA LIKE* (*CSL*) genes were upregulated in *hd1* under LD. Thus, the accumulation profiles of (1-3; 1-4)-beta-glucans and *OsCESA/CSL* transcripts were negatively correlated. This suggests that the differences between wt and *hd1* are mostly due to XyG abundance or that layers of post-transcriptional regulation alter the linear relationship between transcript abundance and cellulose production. For example, interaction between OsCESA subunits forming a functional CSC, transport to the plasma membrane, protein phosphorylation or turnover, may affect the final quantity of cellulose produced.

Among the DEGs, we observed higher expression of some *OsCSL* genes belonging to group C (*OsCSLC*). In Arabidopsis, a quintuple mutant lacking all *AtCSLCs*, could not produce XyG (Kim et al., 2020). This observation suggests that among the *OsCSLCs* upregulated in *hd1*, some might contribute to synthetize XyG, rather than cellulose. Distinguishing between these possibilities will require protein localization studies, since cellulose and XyG biosynthetic enzymes reside on the plasma membrane and on the Golgi membranes, respectively (Cosgrove, 2024). We exclude the possibility of compensatory effects between cellulose and XyG production, i.e. an increase of XyG caused by reduction of cellulose. Such hypothesis has already been tested and discarded by Kim et. al, who showed that plants lacking XyG have normal cellulose content (Kim et al., 2020).

Xylosylation of the glucan backbone is carried out by glycosyltransferases (GTs). We identified several GTs, all of which were upregulated in *hd1* under LD, and partially overlapped with differentially enriched proteins in the proteomic dataset (LOC_Os06g48180; LOC_Os11g18730). These transcriptional profiles are compatible with the hypothesis that *hd1* harbors more ramified XyG in its cell walls. Therefore, Hd1 contributes to cell wall composition in mature leaves. Further studies are needed to understand the implications of this observation on cell wall stiffness and overall plant development.

### Interaction between abiotic stress sensitivity and day length perception

Rice plants are exposed to several abiotic stresses, some of which are exacerbated by climate change, including drought, flooding and exposure to salinity. The latter is particularly relevant in river deltas, where water returning from the sea can intrude for several kilometres in coastal areas. Incorrect water management and fertilizer use also cause soil salinization. Under conditions of high salinity, rice physiology is disturbed by alterations in the osmotic potential, membrane damage, pH instability, as well as the direct toxicity of ions such as Na^+^. Excess salt also reduces photosynthetic efficiency and growth, causes wilting and in severe cases, plant death. The response to salinity is integrated in a global defence system, monitoring environmental and endogenous information, to maximize fitness. Thus, it is unsurprising that part of this system incorporates elements of the photoperiodic response network, which is central in plant adaptation, both in natural and artificial environments. However, how these different pathways communicate with each other remains poorly understood, particularly given the unexpected observation that salinity responses are stabilized across photoperiods by components of the flowering network, and genotypes missing such components, most prominently *Hd1*, respond differently to salt depending on day length.

The closest parallel that we can draw is with drought escape (DE) in Arabidopsis, that is a system better characterized at the molecular level. The DE response allows Arabidopsis to flower early if exposed to water deprivation regimes. This adaptation shortens the life cycle, inducing quick seed set (and paying a trade-off in seed number), if conditions become unfavourable. The trait is photoperiod-dependent, as it occurs under LD, but not SD, conditions. Thus, DE as phenotypic consequence of drought stress, shows differential sensitivity to the photoperiod, similarly to the case of shoot length reported here. Genes within the flowering network, including *GIGANTEA* (*GI*), *FT* and *TWIN SISTER OF FT* (*TSF*) promote the DE response, and plants mutated in these genes flower late, irrespective of watering or photoperiodic conditions (Riboni et al., 2013). This scenario is analogous to rice plants carrying mutations in *Hd1* and exposed to salt, with the notable difference that *Hd1* stabilizes the response in LD and SD, rather than differentiating it.

Drought and other abiotic stresses cause increased production of ABA, in turn coordinating physiological responses, such as stomatal closure, scavenging of reactive oxygen species and osmolyte accumulation (Liu et al., 2022). ABA promotes GI and CO protein activities to induce *FT* transcription in the leaves, resulting in DE (Riboni et al., 2016). Therefore, a plausible interpretation of our findings might be that rice plants exposed to salt stress use Hd1 downstream of ABA signalling to coordinate the responses, in different photoperiods, and that mutations in *Hd1* uncouple day length perception and stress responses. Several lines of evidence support this connection. At the global transcriptional level, it is well established that stress response genes are controlled by the circadian clock, both in Arabidopsis (Covington et al., 2008) and rice (Wei et al., 2022). Altering time measurement by mutating clock genes, prevents proper photoperiodic responses and reduces stress resistance (Wei et al., 2022). Hd1 could be a hub for integration of clock activity and stress responses. If so, more specific hypotheses could be assayed. For example, Arabidopsis PRR proteins, which are integral components of the clock, interact with and stabilize CO during the light phase (Hayama et al., 2017). Triple *prr5 prr7 prr9* mutants have higher tolerance to several stresses, including high salinity, coupled with lower levels of CO protein (Nakamichi et al., 2009). In rice, a clock-dependent mechanism might modify stress sensitivity in the photoperiod, directly as well as indirectly by modifying Hd1 post-translationally. While in Arabidopsis CO protein stability is key to confer a photoperiodic response, Hd1 function is only marginally dependent on its stability, and protein accumulation largely follows transcriptional patterns. Yet, most layers of post-translational protein processing are still to be studied, including phosphorylation or higher order complex formation. We believe the latter could impact on stress tolerance. A homolog of Arabidopsis *PRRs*, *OsPRR73*, is induced by salt stress. If mutated, it increases sensitivity to salt under SD and promotes flowering under LD (Liang et al., 2021; Wei et al., 2021). Importantly, OsPRR73 can form NF-Y complexes, substituting or cooperating with Hd1 to bind DNA. Therefore, Hd1 could be an integrator, downstream of clock-dependent stress responses, or directly involved in controlling expression of stress responsive genes via higher-order complex formation.

## Materials and methods

### Plant material and growth conditions

Rice plants of the Nipponbare and Augusto varieties were used. Augusto carries loss-of-function alleles of *Hd1*, *Ghd7* and *Ghd8*. BC3F3 seeds were obtained by using Augusto as recurrent parental from a Nipponbare x Augusto cross and selecting lines heterozygous for the loss-of-function alleles at each generation. After three rounds of backcrossing, plants were allowed to self-fertilize and in the resulting BC3F2 progeny we selected combinations of homozygous loss-of-function and wild type alleles. The null mutants *hd1-1* and *hd1-2* carry the insertion of a *Tos17* retrotransposon in the first and second exon of *Hd1,* respectively, as described in Gomez-Ariza *et al*. 2015. The *pACT*::*3xFLAG:Hd1* plants were obtained in the Nipponbare background and are described in Eguen *et al*. 2020. The *pHd1:GUS* construct contains the functional Nipponbare *Hd1* promoter and is the same used in Goretti *et al*. 2017.

Plants were grown in phytotrons (Conviron PGR15) at 28°C and 70% relative humidity (RU) during the day and 24°C and 90% RU during the night. Photoperiods were set at 16h light in LD and 10h light in SD. Propagation of plant materials was done in greenhouses at the Botanical Garden Città Studi. Crosses between Nipponbare and Augusto and flowering time experiments of BC3F3 families were done under natural LD field conditions at the Botanical Garden Città Studi in Milan.

### Quantification of mRNA expression and GUS assays

Total RNA was extracted using nucleoZOL (Macherey Nagel) and treated with Turbo DNAse (Thermofisher Scientific) to remove residual DNA. One µg of total RNA was retrotranscribed with the ImProm-II Reverse Transcriptase (Promega) using an oligo dT primer. Quantitative reverse transcription-polymerase chain reaction (qRT-PCR) was used to quantify transcription of individual genes in an Eppendorf Real Plex2.

The list of primers is provided in Supplementary Table 5.

For GUS assays, leaf samples expressing the *pHd1:GUS* construct were fixed in 90% acetone for 20 minutes on ice, kept under vacuum for 1 hour, and then incubated in an X-Gluc solution at 37°C (Jefferson et al., 1987). Subsequently, green tissues were cleared in methanol/acetic acid (3:1, v/v) for 4 hours at room temperature, with constant agitation, followed by multiple washes in 70% ethanol. At least two independent transgenic lines were used, and the experiment was repeated three times with identical results.

### Protein preparation and western blotting

Total proteins were extracted following the protocol described by Wang et al., 2006 (Wang et al., 2006). Proteins were separated by electrophoresis on a 10% acrylamide/bis-acrylamide gel (29:1 ratio). Following electrophoresis, the proteins were transferred to a nitrocellulose membrane and hybridized with anti-Flag overnight at 4°C.

### Chromatin Immunoprecipitation

Chromatin immunoprecipitation was performed using 5 grams of ground tissue powder as previously described, with minor modifications (Perrella et al., 2024). For each experiment, leaves from each genotype were used to extract chromatin. A Bioruptor (Diagenode) was used to shear the chromatin using 40 cycles, each consisting of 30 secs on and 30 secs off, at high power. Anti-Flag magnetic beads (Sigma-Aldrich M8823) were used to immunoprecipitate chromatin. ChIP-qPCR was performed with a 3 min initial denaturation at 95°C followed by 40 cycles at 95°C, 3 secs and 59.5°C, 30 secs. Primers are listed in Supplementary Table 5. Reactions were performed on four technical replicates and three independent biological replicates. Relative enrichment was calculated according to Shapulatov et al. (Shapulatov et al., 2023).

Promoters were defined as genomic regions spanning from −1 Kb upstream, to 100 bp downstream of transcription start sites (TSS) of rice gene models according to Release 7 of the IRGSP annotation of the reference O. sativa Nipponbare genome assembly (Kawahara et al., 2013) (http://rice.uga.edu/). De-novo motif discovery was performed with Weeder 2.0 using the default parameters and conceptual representations of promoter sequences described above. PScan was used to generate P-values for the enrichment of motif PWMs generated by Weeder, scanning the same 1-kb intervals upstream of IRGSP v7 TSSs (Pavesi et al., 2004; Zambelli et al., 2009).

### Quantification of cell wall components

#### Generation of alcohol insoluble residue (AIR)

Leaves of the *hd1-1* mutant and Nipponbare wild type were collected at the same age as for RNA-Seq profiling. The central portion of a mature adult leaf was sampled from 10 plants to produce each biological replicate and ground to powder in liquid nitrogen. The AIR extraction was done following the protocol described by Moore et al 2020 (Moore et al., 2020). Briefly, the powders were made up to 80% using pre-cooled ethanol in 50mL tubes and boiled for 15 minutes to denature any potential cell-modifying enzymes. A destarching step was done with an enzymatic mixture containing amylase and amyloglucosidase from Megazyme (Wicklow, Ireland).

Samples were centrifuged at 2500g for 10 minutes and the supernatant was discarded. Absolute methanol was added to the pellets at 1:10 (w/V) and the tubes were placed on a tube-rotating wheel for 2 hours. After centrifugation at 2500g for 10 minutes the supernatant was discarded. The solvent washing was repeated using equal parts of methanol and chloroform, chloroform, equal parts of chloroform and acetone, and lastly acetone. After the acetone was discarded, excess liquid was allowed to evaporate in the fume hood without letting the pellets dry. The pellets were resuspended in ice-cold deionised water and frozen in liquid nitrogen.

#### Cell wall extraction

The extraction of polymeric material from leaf AIR was performed following a modified protocol from Sathitnaitham et al. (Sathitnaitham et al., 2021). Two extractions were performed using CDTA and NaOH in order to solubilise polymers with varying degrees of association with the cell wall. CDTA was used to solubilise polymers with weak associations to the cell wall (e.g., pectin), while NaOH was used to solubilise more strongly associating polymers (e.g., hemicellulose). For each extraction, 10 mg AIR was weighed into microcentrifuge tubes, and 30 µL of CDTA buffer (50 mM CDTA; 50 mM Tris; pH 7) were added per milligram of AIR. To facilitate efficient mixing, a small stainless-steel ball was introduced into each tube. The tubes were then subjected to mechanical agitation on a Retch Mixer Mill, initially for 2 minutes at 30 Hz, followed by 2 hours at 7 Hz. After extraction, the tubes were centrifuged at 10,000 RPM and the supernatant was stored at −20°C for subsequent analysis. Immediately thereafter, an extraction with NaOH (4 M supplemented with 0.1% NaBH4) was performed, using the same volume and protocol as the preceding CDTA extraction.

#### Enzyme-linked immunosorbent assay (ELISA)

ELISA was conducted on plant cell wall AIR using tissue-culture-treated 96-well plates (Costar 3598, Corning, New York, USA), following the protocol outlined by Sathitnaitham et al. Preliminary tests were conducted to determine an optimal concentration for all rice cell wall extracts to be tested. Appropriately diluted 50 µL aliquots of both CDTA and NaOH extracts were dispensed into the 96-well plates and incubated overnight at 37°C, uncovered. Once the wells were dry, 200 µL blocking agent (3% bovine serum albumin (BSA) in phosphate-buffered saline (PBS)) was added per well, after which the plate was incubated at 37°C for 1h and the BSA/PBS was discarded. Each tested antibody was diluted 1:60 in a solution of 1% BSA in PBS, of which 30 µL was used to probe each well. The samples were incubated at 37°C for 1h and washed 3 times with PBS. A secondary horseradish peroxidase-conjugated antibody corresponding to each primary antibody was diluted 1:10,000 in 1% BSA in PBS, of which 50 µL was added to each well. A final incubation at 37°C for 1h was followed by six washes with PBS. 75 µL of freshly prepared chromogenic substrate (3,3’,5,5’-tetramethylbenzidine at 0.42 mM) was added to each well and the resulting colour-forming reaction was allowed to proceed for 30 minutes before being stopped by the addition of 125 µL sulfuric acid (1 M). Absorbance values were quantified at 450 nm using a Multiskan GO Microplate Spectrophotometer (Thermo Fisher Scientific, Inc., Waltham, MA, USA). Antibodies are listed in Supplementary Table 4.

### Salinity stress assays

The effect of salt stress was analysed in plants grown *in vitro*. Seeds were surface sterilized (one wash in ethanol 70% for 1 minute, followed by 15 minutes of wash in commercial bleach and four washes of 10 minutes each with sterile, distilled water), and placed in culture boxes containing 50 ml of solid growth medium (basal medium Murashige and Skoog with vitamins M0222 Duchefa 4.4 g/L, sucrose 30 g/L, Plant Agar P1001 Duchefa 5 g/L). Ten seeds were germinated in each box. Seven days after germination, 50 ml of liquid medium (same as solid medium, but without Plant Agar) was added in control boxes, and 50 ml of liquid medium supplemented with 300 mM NaCl was added to induce salt stress. Pictures of control and treated plants were taken 14 days after germination and shoot length was measured using ImageJ (https://imagej.net/ij/). The salt sensitivity index was calculated as (shoot length control – shoot length treated)/shoot length control. Control and treated plant pairs were randomly chosen.

### Proteomic analysis

#### Protein extraction

Rice leaves were frozen in liquid nitrogen immediately after sampling, and total proteins were extracted with a pH neutral buffer (4M Urea; 50mM Tris-HCl pH 7.5; 150 mM NaCl; 2mM EDTA; 0,2% 2-mercaptoethanol; 0,1 % SDS; Pierce Protease Inhibitor by Thermo Scientific™). Protein content was quantified by Quick Start Bradford 1x Dye Reagent (Bio-Rad™). Mass spectrometry was performed at the EMBL Proteomics Core Facility in Heidelberg - Germany (https://www.embl.org/groups/proteomics), as described below.

#### Sample preparation

Samples were prepared using the SP3 protocol on a KingFisher APEX system (ThermoFisher Scientific) essentially as described in Leutert et al. (Leutert et al., 2019). Peptides were eluted off the Sera-Mag Speed Beads (GE Healthcare) by tryptic digest (sequencing grade, Promega) in an enzyme to protein ratio 1:50 for overnight digestion at 37°C (50 mM HEPES, pH 8.5; 5 mM Tris(2-carboxyethyl) phosphinhydrochlorid; 20 mM 2-chloroacetamide). Peptides were labelled with Isobaric Label Reagent (Thermo Scientific™) according to the manufacturer’s instructions, combined and desalted on an OASIS® HLB µElution Plate (Waters). Offline high-pH reverse phase fractionation was carried out on an Agilent 1200 Infinity high-performance liquid chromatography system, equipped with a Gemini C18 column (3μm, 110 Å, 100 x 1.0 mm, Phenomenex). Forty-eight fractions were collected and pooled into 12 for MS measurement.

#### LC-MS/MS method

An UltiMate 3000 RSLC nano LC system (Dionex) fitted with a trapping cartridge (µ-Precolumn C18 PepMap 100, 5µm, 300 µm i.d. x 5 mm, 100 Å) and an analytical column (nanoEase™ M/Z HSS T3 column 75 µm x 250 mm C18, 1.8 µm, 100 Å, Waters) was coupled directly to a Fusion Lumos (Thermo Scientific™) mass spectrometer using the Nanospray Flex™ ion source in positive ion mode. Trapping was carried out with a constant flow of 0.05% trifluoroacetic acid at 30 µL. Subsequently, peptides were eluted via the analytical column with a constant flow of solvent A (0.1% formic acid, 3% DMSO in water) at 0.3 µL/min with increasing percentage of solvent B (0.1% formic acid, 3% DMSO in acetonitrile).

The peptides were introduced into the Fusion Lumos via a Pico-Tip Emitter 360 µm OD x 20 µm ID; 10 µm tip (CoAnn Technologies) and an applied spray voltage of 2.4 kV. The capillary temperature was set at 275°C. Full mass scan was acquired with mass range 375-1500 m/z in profile mode in the orbitrap with resolution of 120000.

The filling time was set at maximum of 50 ms with a limitation of 4×105 ions. Data dependent acquisition (DDA) was performed using quadrupole isolation at 0.7 m/z, the resolution of the Orbitrap set to 30000 with a fill time of 94 ms and a limitation of 1×105 ions. A normalized collision energy of 34 was applied. Fixed first mass was set to 110 m/z. MS2 data was acquired in profile mode.

#### MS data analysis

Files were then searched using Fragpipe v20 (protein.tsv files) with MSFragger v3.8 against the Uniprot Oryza sativa japonica database (UP000059680) containing common contaminants and reversed sequences. Contaminants and reverse proteins were filtered out and only proteins that were quantified with at least 2 razor peptides (Razor.Peptides >= 2) were considered for the analysis. The following modifications were included into the search parameters: Carbamidomethyl (C) and TMT18 (K) as fixed modifications, Acetyl (Protein N-term), Oxidation (M) and TMT18 (N-term) as variable modifications. A mass error tolerance of 20 ppm was set for MS1 and MS2 scans. Further parameters were: trypsin as protease with an allowance of maximum two missed cleavages and a minimum peptide length of seven amino acids.

Log2 transformed raw TMT reporter ion intensities (’channel’ columns) were first cleaned for batch effects using the ‘removeBatchEffect’ function of the limma package (Ritchie et al., 2015), and further normalized using the ‘normalizeVSN’ function of the limma package. Missing values were imputed with the ‘knn’ method using the ‘impute’ function of the Msnbase package (Gatto and Lilley, 2012). Proteins were tested for differential expression using a moderated t-test by applying the limma package (’lmFit’ and ‘eBayes’ functions). The replicate information was added as a factor in the design matrix given as an argument to the ‘lmFit’ function of limma. Also, imputed values were given a weight of 0.01 while quantified values were given a weight of 1 in the ‘lmFit’ function. The t-value output of limma for certain statistical comparisons was analyzed with the ‘fdrtool’ function of the fdrtool packages (Strimmer, 2008) to extract p-values and false discovery rates (q-values were used). A protein was annotated as a hit with a false discovery rate (fdr) smaller 0.05 and an absolute fold-change of greater 2 and as a candidate with a fdr below 0.2 and an absolute fold-change of at least 1.5.

### Statistical analysis

Representation factor and p value of overlaps between sets of differentially expressed genes was calculated at http://nemates.org/MA/progs/overlap_stats.html. The total number of genes from the Nipponbare genome was set at 37869 (Sakai et al., 2013). Statistical tests referred to in the text were calculated with Excel or Graph Pad Prism ver.8.0.1.

## Figure legends

**Supplementary Figure 1.** RNA sequencing controls and phase enrichment of DE genes. **A**, flowering time of Nipponbare wild type and *hd1-1* mutants under LD (blue) and SD (green). Each symbol corresponds to one plant. The number of plants scored is indicated in the histograms. **B-C**, expression of *Hd3a* and *RFT1* under LD (**B**) and SD (**C**) in the samples used for RNA sequencing. **D**, results of phase enrichment analysis obtained using Phaser. The number of genes is plotted against the phase of expression. ZT, *Zeitgeber*.

**Supplementary Figure 2.** Overlap between genes controlled by *Hd1* and the photoperiod. The Venn diagram shows the intersection between genes regulated by *Hd1* under LD (*hd1* LD), SD (*hd1* SD) and the shift from LD to SD (LD to SD). The size of the circles is proportional to the number of differentially expressed genes. The LD to SD dataset is from Galbiati *et al*., 2016. Genes were filtered for FDR≤0.01 and logFC≥|1.5|.

**Supplementary Figure 3.** Gene ontology categories enriched in the *hd1* transcriptomes. The graph shows categories significantly enriched more than 1.5 folds under LD and SD. Enrichments were performed using release 2024-01-17 of the Gene Ontology Resource database, with default settings. Input gene lists were filtered for FDR≤0.05 and log_2_FC≥|1.5|. Note that due to the small size of the SD dataset, few genes can cause a very high overrepresentation of some terms. Except for inflorescence development and GA mediated signaling pathway, categories are represented by *SPX1* and *SPX2* only, both of which are downregulated and control several developmental and physiological processes (Wang et al., 2014). CC, cellular compartment; MF, molecular function; BP, biological process.

**Supplementary Figure 4.** Genes regulated by *Hd1* in the phenylpropanoid biosynthetic pathway. The image shows a scheme of the phenylpropanoid pathway as retrieved from the Kyoto Encyclopedia of Genes and Genomes (KEGG) database. The coumarine branch of the pathway was added manually. Gene identifiers are either red or blue when upregulated or downregulated under LD, respectively. Note that enzymes are often encoded by multiple genes.

**Supplementary Figure 5.** The *hd1* leaf proteome under LD and SD. **A-D**, volcano plots showing differentially abundant proteins in the indicated comparisons. Expression coordinates are determined by -log_2_FC and -log_10_(p-value). Red indicates protein hits (fdr<0.05 and absolute fold-change>2); blue indicates protein candidates (fdr<0.2 and absolute fold-change>1.5); green indicates proteins not significantly different between genotypes or treatments. Note that differences are mostly detected between photoperiods.

**Supplementary Figure 6.** Ontological categories of differentially expressed proteins. The graph shows gene ontology categories enriched in the lists of differentially abundant proteins, in the indicated comparisons. The size of the circles indicates the extent of enrichment; the color indicates the adjusted p-value.

## Acknowledgements

We are grateful to the personnel of the Botanical Garden ‘Città Studi’ for support with plant care, and to Jennifer Schwartz and Frank Stein of the EMBLEM Proteomics core facility for support with proteomic analyses.

## Author contributions

F.F. designed the study and wrote the manuscript. M.B., D.C., A.B., E.B., F.D., E.F. and T.E. collected samples, performed molecular work and analyzed data. B.K., D.H. and M.C. performed the bioinformatics analysis. F.T., J.P.M., E.T., S.W., F.L.S., I.E., V.B., G.P. and C.B. interpreted the data and revised the manuscript. All authors read and approved the final manuscript.

